# Effect of microampere-scale wireless conductive microelectrostimulation on *Aspergillus fumigatus* growth on solid cultures

**DOI:** 10.64898/2026.08.10.743807

**Authors:** Manousos E Kambouris, Stavroula Kritikou, Afroditi Milioni, Gian Marco Ludovici, Katerina Karageorgou, Aristea Velegraki

## Abstract

The effect of microcurrents on facultative microbial pathogens remains controversial. Solid cultures in Sabouraud Glucose Agar of the ubiquitous mold *Aspergillus fumigatus* were repeatedly treated with a commercially available device performing wireless conductive microelectrostimulation by 3.5 μA microcurrent routed by spraying negatively charged air particles onto solid cultures in modified petri dishes. The treated cultures displayed increased growth compared to standard ones, but only as a function of mycelial density and total surface; the radial growth rate of the mycelium remained unaltered. The increased growth was positively related to the duration of the treatment. At the same time, secondary development (new mycelial loci within the dish) was greatly upheld due to treatment, as the spraying created microairstreams dislocating the fungal spores. These results imply perplexed kinetics of mycelial growth both with and without treatment, since the folding of the mycelial mat is observed regularly. Both the fungus’ response to the ES and the possible revision of growth kinetics create prospects for biotechnological and bioremediation applications but also imply biomedical considerations, regarding infection dynamics of mycelial fungi and their *in situ* resistance to immune responses and treatment.

## Introduction

Channeling electric current through multicellular organisms has long been a known procedure with various results. High intensities cause electrocution while lower intensities are used for electrotherapy or for enhancement of physical training. Studying the effect on microorganisms of various forms of electricity, such as electric fields (EFs) or different types of electric current, gains momentum, due to the possible applications. The manipulation of the microbial growth may lead to new approaches in waste disposal and decomposition, in biomass and secondary metabolite production, in allogenic *in vivo* expression and to many other applications ^1–3^. The antibiotic effect of electric fields is documented since the 60s ^4^ for Alternate Current (AC) fields; Direct Current (DC) pulsed fields were successfully studied in the 90s ^5^, while static fields furnished inconclusive results ^6^, as do magnetic and electromagnetic fields ^7^. Actual current transfer, though, is another issue altogether. There are many studies, differing in the type of current, may that be AC, DC or other ^8–10^ or the type, design and polarity of the electrodes used ^11^ and, of course, dosage ^12^. A number of studies describe positive ^1,3,13^ and another, considerable number, negative effects in microbial growth ^1,6,8–10^. The matter clearly remains open and undecided ^2,7^, but the bulk of the published work is performed using contact electrodes onto unicellular organisms; mostly bacteria, but occasionally some yeasts as well ^13^.

Reasons for inconsistent results may be the different individual tolerances of microbes and the differences in electricity quantity and quality. The quantitative parameter may be differentiated to one electrodynamic attribute, referring to the cumulative electric charge supplied to the organism/cell, and to one electrokinetic attribute; the latter refers to the rate of supplying the given charge (which is described by the intensity and, when applicable, by the frequency and the waveforms). The qualitative parameter is more heterogeneous and comprises various attributes such as the material, design and interface of electrodes, the circuit architecture and the use of EF as opposed to the use of electric current.

Wireless Conductive Microelectrostimulation (WCMES) emerged in early 2010s. It refers to ionizing atmospheric gases (either Oxygen or Nitrogen), condensing them onto airborne water molecules and spraying the produced ionized conglomerates (negatively/positively charged, respectively) onto the target surface, which is placed on an insulated base or pedestal; the target is connected by an adjustable neutral electrode, which remains outside of the spraying footprint, back to the device to close the circuit ^14^. The W200 used herein sprays ionized Oxygen, of negative ion value, and is effective at a distance of less than 12 cm, while the output intensity of the microcurrent is 0.5-4 μA at 0.5 μA intervals.

A robust answer on whether the WCMES has a positive or negative effect on microbial growth is of paramount importance for biotechnological applications, as non-medical cultures could be affected in a desirable way; the increase of yield, either biomass or metabolites, and the reduction of harvesting time or the decrease on incubation temperatures being especially lucrative perspectives. Alternatively, negative effects might imply potential for sterilizing or decontaminating use, bypassing adverse effects of ionizing irradiation and environmental chemical/pharmaceutical burden.

In this study, extremely low equivalent intensity of μA level is used for WCMES treatment of *Aspergillus fumigatus* cultures. As a eukaryotic, multicellular, facultative pathogenic mold of extremely wide geographical distribution and of high prevalence in disease environments ^15^, *A. fumigatus* is a suitable model for studying a more complicated lifeform with biotechnological, environmental and biomedical impact. The universal distribution of *A*. *fumigatus* is a key factor in respiratory conditions, both allergies and infection ^16–20^ and results in colonization of exposed human biocompartments.

## Materials and Methods

### Strains & Culture conditions

Two clinical strains of *A. fumigatus* (HCPF-8165 and HCPF-14960) from the Hellenic Collection of Pathogenic Fungi were used for different purposes (pilot and main experiments). Sabouraud Glucose 2% Agar with Chloramphenicol (SGA- Carl Roth GmbH + Co. KG, Karlsruhe, Germany), was prepared as suggested by the manufacturer; the ready- to - dissolve SGA powder was enhanced with 0,3% w/v Agar to improve sturdiness and, after autoclaving according to the manufacturer, was cast into sterile plastic 9-cm petri dishes under aseptic conditions.

The organism was puncture-inoculated in three duplicate Sabouraud Agar petri dishes. One dish per pair was modified to become conductive, while the other was of standard form and use. Conductive dishes were produced by embedding separately autoclaved aluminum (Al) foil strips at the petri dishes just prior to casting the medium so as to fold the two edges of the Al strip over the dish lip and down under the dish underside. The dish was placed on a metal plate, which was thus in contact with the folded parts of the aluminum strip and also collected the ion flow sprayed outside the perimeter of the dish, thus increasing the return of the sprayed charge back to the device. The whole was seated on a flat, rectangular piece of Expanded Polystyrene acting as insulating material (**Figure** 1A-B). Thus the strip closed the circuit with the ions transferred through the air from the head of the device. A second version of conductive dishes was produced with Al foil strips only at the bottom, and the underside of the dish pierced by standard copper pins at the footprint of the foil (**Figure** 1C).

**Figure 1.**
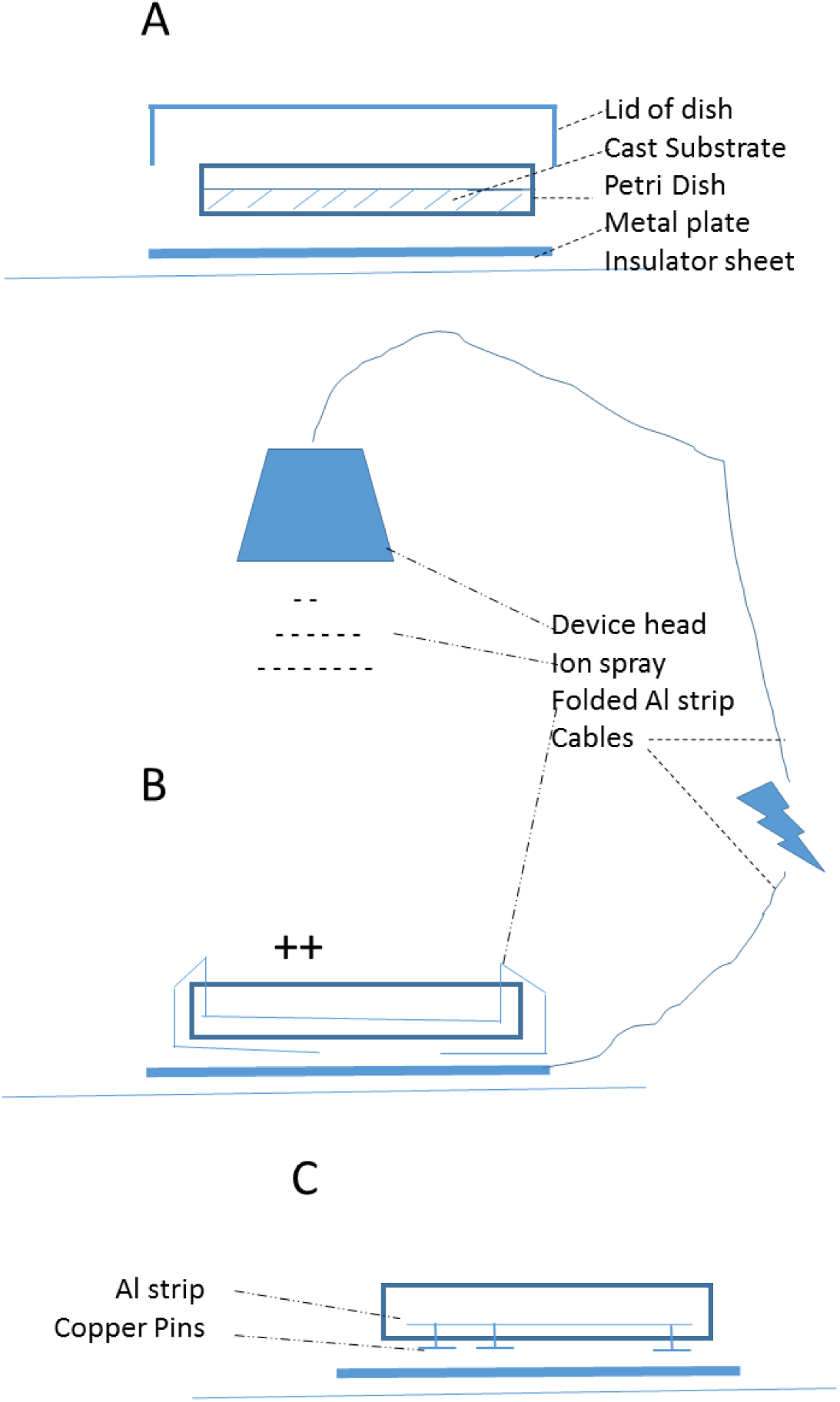
Conductive Petri dishes and treatment. A: The setup of the cast petri dish on the metal plate and sheet of Expanded Polystyrene for insulation B: The setup of treatment of the conductive petri dish with expanded Al foil strip, bent downwards, by the W200 Oxygen-spraying device. The nutrient substrate is not shown for simplicity but it covers the Al strip at the bottom of the dish. C: The setup of treatment of the conductive petri dish with short Al foil strip, and the bottom pierced by copper pins functioning as conductive bridges. The nutrient substrate is not shown for simplicity but it covers the Al strip at the bottom of the dish.

Three conductive dishes have been subjected to 3.5 μΑ Wetling (Wetling, Danemark) Oxygen-ionizing device treatment for 1, 10 and 60 min respectively, while placed on a metal plate, itself lying on a non-conductive piece of polymer (Expanded Polystyrene) after being puncture-inoculated with strain HCPF-14960. Treatment sessions were performed daily, except for Day 2. The Standard dishes were left open concurrently to the WCMES treatment same time, on the same piece of Expanded Polystyrene, but outside the device’s spraying footprint and off the metal plate. The dishes were incubated at room temperature. Every day the developing mycelia were measured; at first the diameter (2r) was measured by ruler, as the maximal distance between two opposite points of the periphery, and after the mycelia acquired a minimum size, the measurement was performed from the puncture point to the furthest outboard point, counting radius (r) rather than diameter. Measurement of the maximal radius is standard practice whenever mycelia are not circular. Human error must be taken into account, especially as the active edge (peripheral growth zone) of the mycelium is colorless and diathlastic. Consequently, a certain arbitrary observation angle was needed for it to become visible. But observation under such angle impairs accurate measurement with the ruler.

Secondary mycelia, generated from spores of the primary mycelium displaced by the air stream of the spraying or by environmental contaminants were enumerated but not measured. For the first three days the accumulative Dosage (in μCb), as given by the instrument after the conclusion of each treatment session, was noted, as an indicator of variability, especially among supposedly identical treatment sessions of the same dish in successive days.

In a separate experiment, the HCPF-14960 strain was puncture-inoculated into one dish with aluminum foil strips and one standard dish, both which were subsequently incubated at room temperature without any WCMES treatment, to evaluate any possible interference of the presence of the aluminum foil *per se,* when inserted in the substrate, with the fungal growth. These results would set the baseline comparison for the main experiments.

Moreover, a pilot experiment was conducted with the HCPF-8165 strain. It was puncture-inoculated to one standard petri and three conductive (sample) ones, each of the latter exposed to one session of treatment, right after inoculation: one Petri exposed for 1 min, one for 10 min, and one for 1h. Thus, the standard Petri dish remained open, to emulate the conditions of the samples, for 71 min, which makes a very important difference in terms of free aeration time.

## Results

The HCPF-14960 inoculated in the untreated conductive petri dish implanted with aluminum-foil strip (Al+) exhibited markedly less growth than the respective standard petri dish (Al-), with a diameter of 4 mm compared to 6 mm for the first day and 24 mm to 28 mm for the second day. After 7 days, the mycelial mat in the conductive dish was thin, whereas the mycelium mat of the standard dish was dense and with radial folding (**Figure** 2). This suggested a markedly adverse effect caused by the aluminum-foil strip embedded into the solid medium of the culture compared to its respective standard, amounting to 50% the first day, and 17% the second day as a function of the mycelial diameter (**Table** 1).

**Figure 2.**
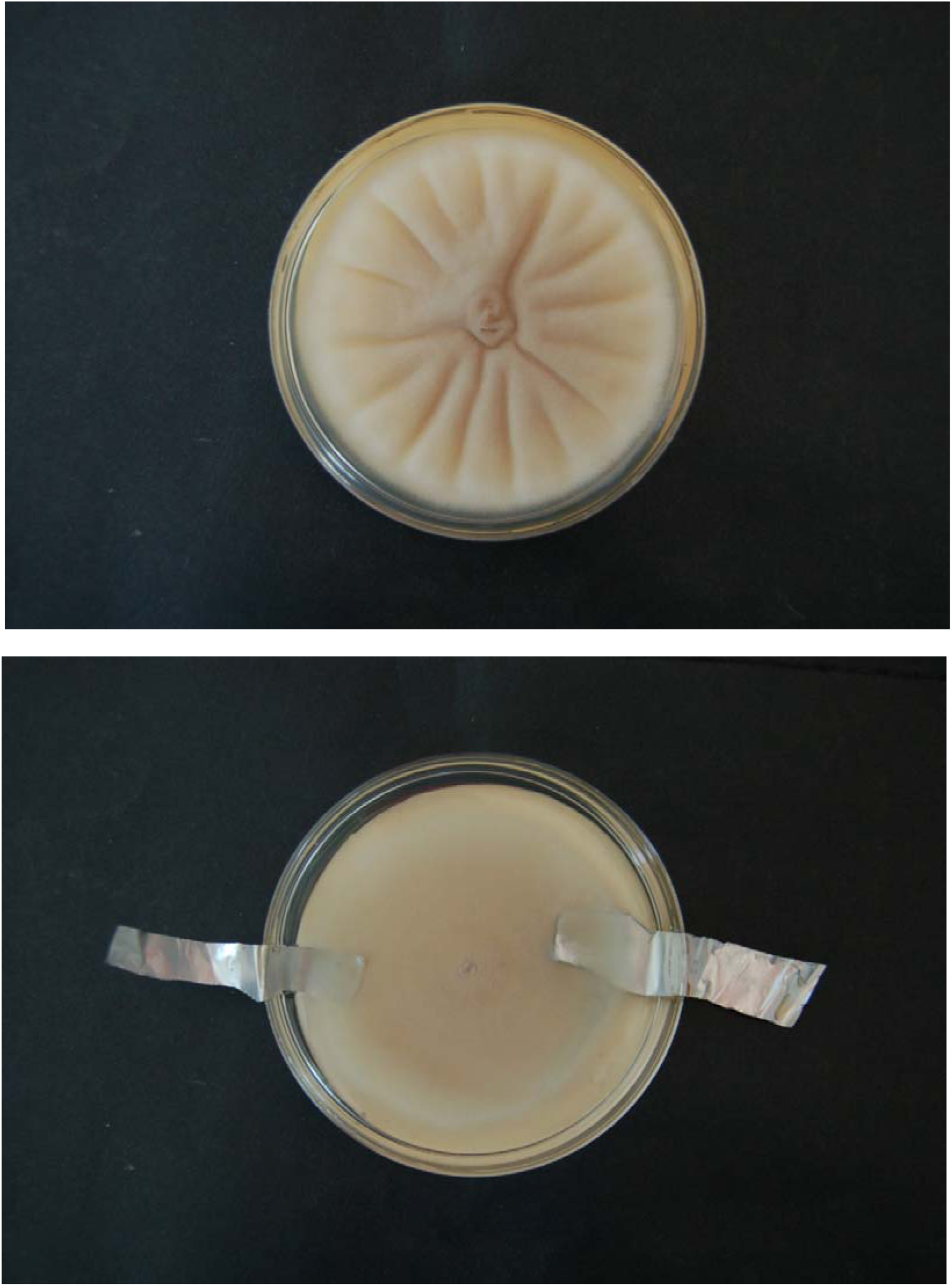
Net effect of Al in mycelial growth. Non-exposed cultures with (bottom) and without (top) embedded Aluminum foil strips (Al) at their bottom, incubated concurrently. Everything else being identical, the presence of Aluminum foil decreases growth as a function of density and lateral expansion, producing no radial folds

**Table 1.**
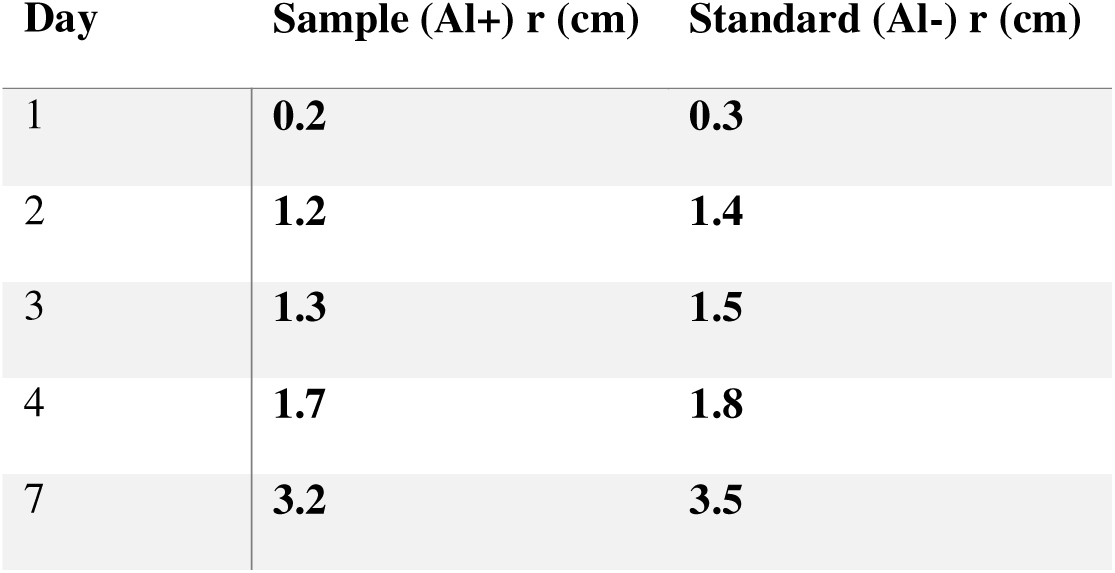
No WCMES, comparative growth (r) of HCPF 14960 in the presence (+)/absence (-) of Aluminum foil (Al) strip. Aluminum foil implants affect negatively the mycelial growth.

| Day | Sample (Al+) r (cm) | Standard (Al-) r (cm) |
| --- | --- | --- |
| 1 | 0.2 | 0.3 |
| 2 | 1.2 | 1.4 |
| 3 | 1.3 | 1.5 |
| 4 | 1.7 | 1.8 |
| 7 | 3.2 | 3.5 |

Pilot experiments of single-dose WCMES treatments administered immediately after puncture inoculation (strain HCPF-8165) compared against a single untreated standard dish indicated a faster growth for the mycelium treated for 1h by WCMES - compared to the standard. The sample treated for 1min failed to grow altogether; and the sample treated for 10 min grew markedly less than the untreated standard (**Table** 2 & **Figure** 3).

**Figure 3.**
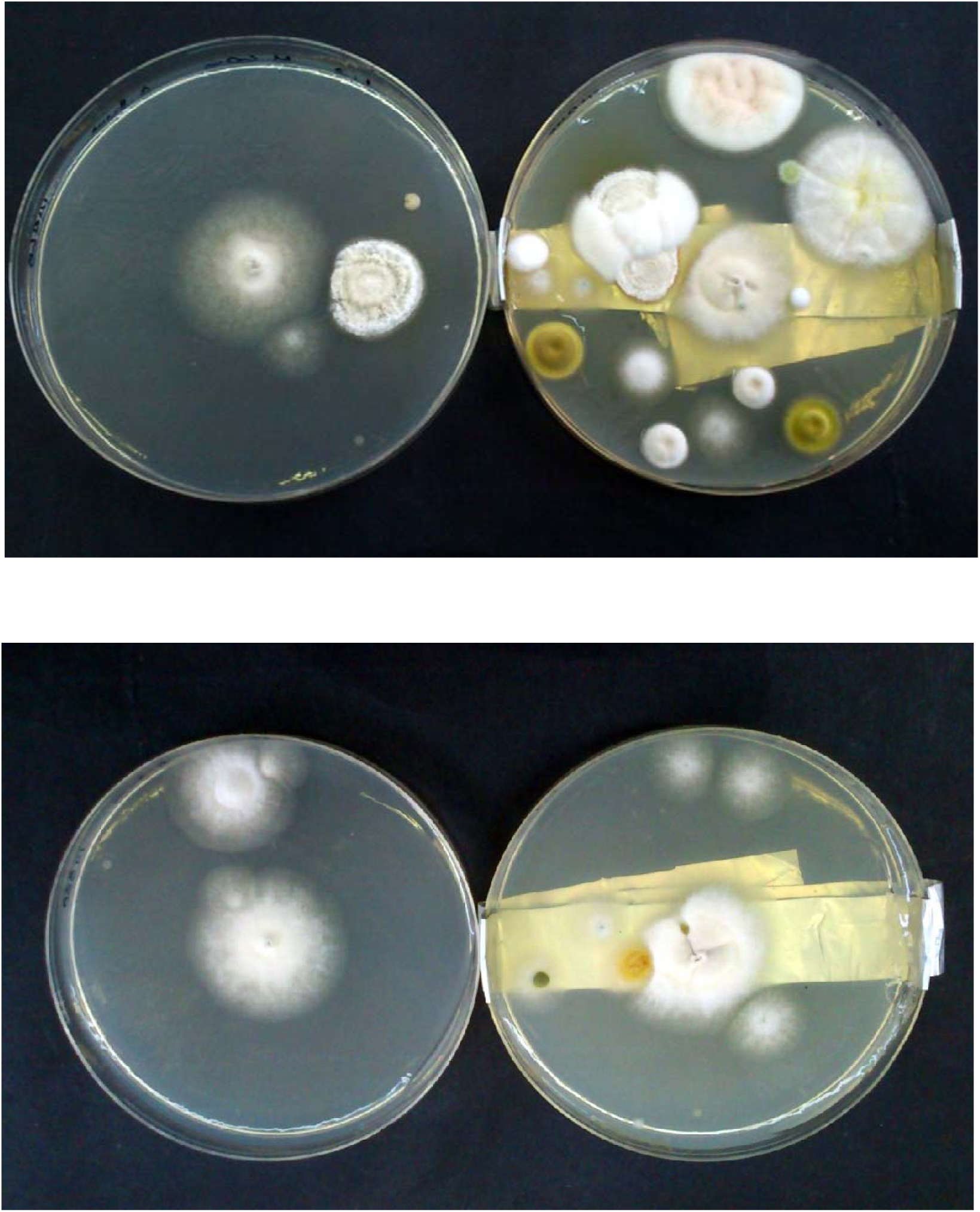
WCMES effect on secondary growth and contamination increase. Secondary growth increases with treatment (left: untreated, right: treated) and treatment time (upper photo: 1h; lower photo: 10min) even in cases receiving treatment only once.

**Table 2.** WCMES dosage and mycelial growth of once-treated puncture cultures of *A. fumigatus* HCPF-8165 on SGA.

| Dish | Ion Dosage (μCb) | 2r<sub>d1</sub> (cm) | 2r<sub>d2</sub> (cm) | 2r<sub>d3</sub> (cm) | 2r<sub>d4</sub> (cm) | 2r<sub>d5</sub> (cm) |
| --- | --- | --- | --- | --- | --- | --- |
| Standard | — | 0.6 | 1.4 | 2.4 | 3 | 4 |
| Sample 1 min | -185 | — | — | — | — | — |
| Sample 10 min | -2,045 | 0.2 | 1 | 2 | 2.8 | 3.2 |
| Sample 60 min | -11,121 | 1 | 1.8 | 2.6 | 3 | 4.6 |

All three repeatedly treated dishes (HCPF-14960) had similar -although not identical- radial growth rates of their mycelia for the duration of the experiment (see **Tables** 3-5). The results suggest mostly kinetic rather than dynamic differentiation in the radial extension of the repeatedly treated mycelia, compared to respective standard dishes.

**Table 3.** WCMES Dosage, radial mycelial growth and secondary loci of A. fumigatus HCPF-14960 puncture culture on SGA treated daily for 60 min.

| Day | Ion Dosage<br>($\mu\text{Cb}$) | Standard<br>2r (cm) | Sample<br>2r (cm) | Standard<br>CFU | Sample<br>CFU |
| --- | --- | --- | --- | --- | --- |
| 0 | -15,101 |  |  |  |  |
| 2 | -12,523 | 0.4 | 0.6 |  |  |
| 3 | -12,518 | 1.2 | 1.2 |  |  |
| 4 | -12,662 | 1.6 | 1.6 | — |  |
| 5 |  | 1.8 | 1.8 |  | — |
| 6 |  | 2.4 | 2.2 | 4 | 12 |
| 7 |  | 3.2 | 3.2 | 4 | 17 |

**Table 4.** WCMES Dosage, radial mycelial growth and secondary loci of *A. fumigatus*

| Day | Ion Dosage<br>($\mu\text{Cb}$) | Standard<br>2r (cm) | Sample<br>2r (cm) | Standard<br>CFU | Sample<br>CFU |
| --- | --- | --- | --- | --- | --- |
| 0 | -2,345 |  |  |  |  |
| 2 | -2,121 | 0.6 | 0.7 |  |  |
| 3 | -2,121 | 1.2 | 1.2 |  |  |
| 4 | -2,068 | 1.8 | 1.8 | — |  |
| 5 |  | 2.4 | 2.4 |  | — |
| 6 |  | 2.8 | 2.4 | 4 | 8 |
| 7 |  | 3 | 2.8 | 4 | 8 |

**Table 5.** WCMES Dosage, radial mycelial growth and secondary loci of *A. fumigatus* HCPF-14960 puncture culture on SGA treated daily for 1 min.

| Day | Ion Dosage<br>($\mu\text{Cb}$) | Standard<br>2r (cm) | Sample<br>2r (cm) | Standard<br>CFU | Sample<br>CFU |
| --- | --- | --- | --- | --- | --- |
| 0 | -212 |  |  |  |  |
| 2 | -206 | 0.4 | 0.4 |  |  |
| 3 | -213 | 0.8 | 0.8 |  |  |
| 4 | -207 | 1.4 | 1.4 | — |  |
| 5 |  | 2.4 | 2.4 |  | — |
| 6 |  | 2.6 | 3 | 5 | 3 |
| 7 |  | 3 | 3 | 5 | 3 |

The treatment sessions had to be discontinued at day 5 as secondary autologous and xenologous mycelia were plentiful and visibly interfering with the primary mycelial growth (**Figures** 3,4). For the same reason, diameter/radius measurements were performed only up to Day 8. Contaminations (xenologous growth) and secondary mycelia (autologous growth) were plentiful, the time of exposure being as important a bolstering factor as the exposure proper. The mycelia count in the 60min-sample was 212% higher than in the 10min-sample and 566% than in the 1min-sample (**Tables** 3-5). Similarly, for 10min- and 60min treatment, samples yielded significantly higher mycelial count than standards; 100% and 425% respectively (**Figure** 4 & **Tables** 3-4).

**Figure 4.**
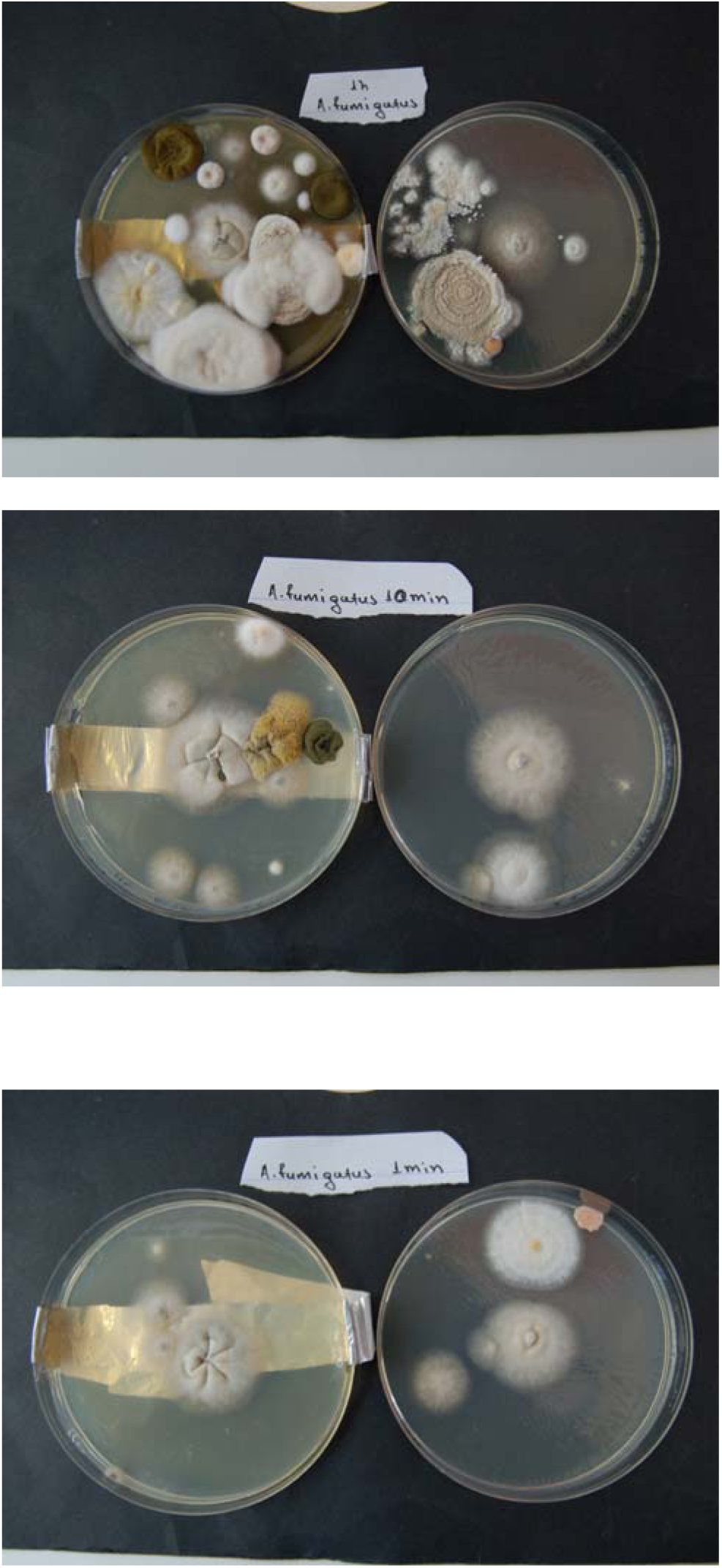
WCMES effect on mycelia growth. Treated and untreated plates exhibit difference in mycelia texture. The former exhibit more thickly woven and folded mycelial mats, which become more prominently so as treatment time increases.

The most intriguing observation was the obvious difference in the texture of the mycelia. Untreated mycelia were thin-woven with sparse hyphae while the 60 min- repeatedly treated mycelium was thick and dense; the other two repeatedly treated mycelia, exposed for much less time (1 min and 10 min) were clearly less dense upon conclusion of the experiment (Day 7). Dishes were kept for another 10 days at RT, and were photographed before being discarded. All treated dishes had developed folds in the primary mycelium, the depth of which was a function of the treatment time; the thin-woven untreated mycelia developed no folds whatsoever. All mycelia at the moment of this observation had been of the same apparent diameter; "apparent" is underlined, since non-radial folds equal to increased true diameter (**Figure** 4).

## Discussion

### Growth intricacies and conditions

The adverse effect of the embedded Aluminum foil strip (**Table** 1 & **Figure** 2) clearly suggests that any increase in growth (radial or, more probably, lateral) that would be observed in the WCMES-treated dishes compared to the respective standards was to be attributed to the WCMES proper, the positive effect of which is the opposite, actually antagonistic, of the negative effect of the presence of Aluminum foil.

Regarding the comparative results of the dishes treated only once (HPCF- 8165) there are two factors that may explain the unexpected comparison of the Standard with the samples exposed for 1 and 10 min (**Table 2** & **Figure** 3). The one is the difference of the availability of fresh air, since the 10min sample was left open for 10min, under treatment, while the standard was left open for 71 min. The other, not exclusive to the just mentioned one, is the negative effect of the Aluminum foil strip in the growth, as substantiated above. The WCMES treatment for 1 min and 10 min was too feeble for its inductive effect to overcome the deleterious effect of Aluminum foil; thus the sample treated for 1 min failed to grow any mycelium, while the sample treated for 10 min was induced enough to produce growth, but not enough so as to overcome the handicap of the presence of Aluminum foil and thus the growth is less than that of the standard. The 60-min treatment was enough to overcome the handicap, though, and produce increased growth compared to the standard. The accumulative dosages give an idea of the comparative induction exercised upon the mycelia. Similarly scale results can be seen in the dishes receiving repeated treatment (**Tables** 3-5), although the aggregated nature of the treatment would be expected to produce, even in short treatments, a considerable accumulated dosage and thus effect.

### The medicinal dimension

The effect of electrical modalities on the growth of microbiota has been previously shown to be highly conditional; the same settings affect differently different organisms, especially different mycelial fungi ^1^, which implies that for the same effect, different settings should be devised, obviously after testing- a concept defining the field of Electroculturomics ^21^. This seems applicable mostly in a biotechnological context. The direct medical implication is that such treatment, for whatever ailment, is ill-advised when an active infection is detected or suspected ^2,22^. The comparative effects on hosts immunity mechanisms, on the physiology of the pathogen, and on that of the host are impossible to predict and even if simulated with high fidelity through Big Data approaches ^23–25^, physiology particularities among hosts would make generalized guidelines inadvisable, despite hard evidence of antimicrobial effect *in vivo-in situ* ^22,26–29^. However, the microbial presence in biocompartments previously considered sterile ^30^ and the intrinsic associations of microbiota within appendage microbiomes ^31–33^ also carry a degree of uncertainty. Still, the decades of use of diverse Electroceuticals for a range of interventions ^34,35^, from pain management ^36^ to wound healing ^37^ and psychiatrics ^38^, allow optimism on the matter of Electroceuticals in widespread therapeutic use.

The most important connotation of this work is not in the field of prospective applications, though. The kinetic- *not* dynamic- differentiation in the radial growth, as seen in Tables 3-5, may affect the response of the host’s immunity. First due to faster Vs slower initial radial speed of growth and thus dissemination potential. Then, by the faster Vs slower exposure of response-evoking antigenic markers. The most important issue though is the preferential lateral rather than radial growth, as it is evident from the comparison of the **Figures** 2-4 with the respective **Tables** 1-5. The Figures show the density and lateral growth as a visual, semi-quantitative marker; the Tables show quantitatively the radial growth. These observations further validate previous conclusions on mycelial dynamics and kinetics under duress ^39^. This preferentiality probably occurs in increments, under such –or any- induction, as suggestive by the concentric rings of different colors, wherever/whenever these appear, in growing mycelia, depending on species and incubation conditions.

A close-knit, dense, slowly advancing mycelium rather than an aggressively expanding but thinner one in a respiratory, or other, infection, is of grave importance. The mycelial status itself protects the fungal cells from phagocytosis due to size through structural continuity, leaving only molecular amenities, such as (but not limited to) the attack complex of the Complement as a practical antifungal mechanism. A close-knit conformation would restrict, if not negate altogether, the access of molecular agents to a large proportion of the mycelial cells. Amongst these are the Complement and antibodies, but also other active, micro-molecular moieties. The aspergilloma may be a good example on the issue, which is not expanding in principle, but may be calcified and thus highly resistant to rupture mechanisms ^17,18,40^. Such dense packing of hyphae would also allow faster repair and/or replacement of necrosed mycelial cells, at least while the mycelium is young ^39^.

In the longer term, manipulation by electrostimulation could limit the invasive phase of an infection by encouraging lateral growth, and once it is contained, it might re-initiate axial growth of the hyphae, to allow better access of antifungal agents to the mycelium, and to discourage the production if toxins and other environment-modifying agents, including lytic enzymes, the optimal production of which is the idiophase, occurring after the trophophase (expansive phase of growth), as known from industrial batch cultures ^41,42^.

WCMES and colonization.

Contamination and secondary mycelia development were obvious and related more to treatment conditions and less to simple exposure of the dishes, staying without lid in indoors space (bench), as obvious from the relatively steady mycelial count in untreated dishes. This is due to the air stream produced by the W200 device for the spray of the ions, which favors the passive flight of fungal conidia. The difference in counts of Mycelium Forming Units/MFUs between dishes treated for 60 min and 10 min (**Tables** 3 and 4) is indicative of the said effect of the airstream on the primary mycelium. The relation, although causative, is not proportional; this is readily understood, as within some minutes all ready- to-fly spores become airborne. The rest are not ready to detach, nor are they prone to be. The airstream, thus, even if steady, produces a peak of spore delivery and not a continuous flow. Thus, the aerodynamics alone suffice for explanation of time/MFU counts disproportionality making unnecessary any implication of the electrodynamic effect of the device.

The creation of secondary mycelia (defined as new mycelia derived from MFUs, i.e. spores or other detachable bodies grown on the originally inoculated mycelium/colony) is a very important factor in the total increase of biomass, but incorporating it into growth modeling is difficult, especially so for sporulating fungi. Such an increase would be important in both biotechnological and medical contexts. The growth of contaminating mycelia and airborne colonies may be lethal when dealing with open wounds, like ulcers and thermal or chemical burns ^14,34,43,44^ as they pose an equal danger for (re)infection and dissemination. In biotechnological applications it becomes far more alarming, as it may compromise cultures of high market values.

### WCMES and mycelium kinetics

The texture of the repetitively/daily treated mycelia compared to the respective standards clearly indicates an enhanced growth, implemented laterally instead of axially, causing neighbor hyphae to shove, fuse and push, thus producing a 3-D development which cannot be currently seen due to the lack of a rigid fungal framework to support perpendicular development. Had it not been so, it is possible that a spherical surface, a dome, would emerge, its apex (vertical radius) being proportional to the depth and number of mycelial folds, i.e. fungal growth observed. The unfolded mycelial mats would then produce flat patterns, the slightly folded ones would produce a slightly convex and the intensely folded ones intensely convex dome-like (or tubular ring-like) structures.

### WCMES and mycelium dynamics

It is a well-established fact that the mycelial growth pattern, entailing hyphal branching, favors lateral rather than axial growth ^45^ due to the need of the fungus to optimally use biomass for the exploitation of a given substrate area ^46,47^. The distribution of nutrients is unlikely to follow an axial pattern due to dispersion, diffusion and entropy principles. Apical branching (split-tip branching) tends to favor extension rather than expansion ^48^ in order to increase the area coverage of the mycelium and balance the double task of exploring as far as possible and exploit resources as widely as possible ^47^. This double function is best served by a circular extension pattern, and circularization principles explored previously ^49^, are felt to be part of the switching of the tips from mostly apical to mostly lateral branching. If branching angles are high, 45^0^ or more, the expansion is favored but this creates peculiar multi-focal colonies with a tendency to regress to the germination point and creating a hive-like pattern. Split-tip branching tends to maintain the angular density of hyphae over the solid substrate, possibly to a balance where the available substrate may support a growth ratio inducing a two-pronged (which is double, in energy currency) development ^48,50^. Although a negative autotropism mechanism has been suggested ^51^ to establish a minimum of separation between the splits of a hypha, the maximum separation might be under the control of a "persistence factor" ^52^. It is plausible that the actual control mechanism and the causative basis of triggering and directing apical branching may not coincide, as the two abovementioned mechanisms may co-exist to provide a highly accurate control mechanism.

Such concerns do not favor 3-D branching or ascending growth, as the air has preciously little growth factors to provide to a terrestrial organism -but for the oxygen. This is an excellent reason for the evolutionary lack of any supporting framework in fungi and in sharp contrast to Plantae, where positioning for light reception is of premium importance. But once volume becomes of consequence for mycelial fungi, a basic 3-D diffusion/ accommodation of the extra growth efficiency is achieved by folding. Thus, the symmetrical, usually radial folding observed in many puncture cultures may not be a defining character (as is sclerotial and droplet formation, number and positioning). It may rather be an adaptational, reactive feature, the intensity of which depends on relative axial and lateral growth rates conditional to external factors such as incubation parameters and substrate composition. If such is the case, the observation of neat, symmetric radial folds might be indicative of reaching a given growth level in the development of a mycelium.

Pushing the issue further, the presence of more than one ring of such radial folds, of different fold number and size, might imply different and periodic intensive growth cycles, much to the like of tree trunk rings. Such a thought might have been implied previously ^47^ when suggesting that rules governing branching patterns may change during the lifetime of a mycelium. Herein, the idea of periodic change is added.

On the other hand, the radial change of color in mycelia is not yet linked to specific physiological changes in growth dynamics. Obviously, the growth dynamics of moulds should be reconsidered, as such pulsed dynamics and 3-D branching potential could differentiate their growth patterns within a substrate, might it be an infected individual or a decomposing, valuable or harmful mass of dead matter, such as artefacts and organic waste respectively. Up to now 3D modelling ^53^ refers to the dynamics within a branching or elongating hypha and not to different hyphae expanding in space rather than surface. This restrictiveness in perception is understandable, since the 2-D growth pattern is deemed one key differentiation character of fungi towards animals and plants^54^. +Whether and how the WCMES affects and/or is affected by the 3 phases of hyphal growth ^51,55^, i.e. whether it extends the exponential one or triggers some transition from one to another, is yet unresolved. Obviously, the kinetics and the direction of such transition or transitions (induction/retardation) or some switch from axial to lateral growth and vice versa remain matters of conjecture. Resolving them may require some quantitative biomarker, like the mathematical description of apical elongation and branching events on one hand and a 2-D, surface density computation on the other. The folding of the mycelial mat implies a possible need for 3-D computation, maybe not based on volume density but on surface density of 3-D shapes, functioning as surface projection of volume differentiations.

### WCMES in yeast: a matter of perspective

The budding of yeasts is in effect a branching process with less suppression enacted. Multiple "branching" events of one step each, following a much more liberal 3-D pattern in spatial terms but never coexisting in the same temporal frame might be a rather accurate description of budding using branching terminology. Some yeasts form pseudohyphae and the dimorphic fungi have both yeast and true hyphae phases ^56^. Moreover, *Schizosacharomyces* peculiar breeding (by medial fission rather than by budding) is halfway the evolutionary distance between yeast cell and mycelium thallic systems, as it combines the daughter cell separation of the former with the spatial self-restriction of the latter ^57^. With all that in mind, observations of both thallic systems should be viewed preferentially through *sensu lato* criteria. Any additional growth in yeast population through MES does not need to be differing from what has been observed and described for mycelial fungi/forms.

### WCMES in a wider context

Projections of WCMES application to other lifeforms like bacteria, Animalia and Plantae may be of relevance. The electrostimulation has been found to increase extracellular matrix production such as elastin and collagen in human cells ^58^, thus increasing healing rate by enhancing repopulation and repair. Electronegativity, in particular, seems to be especially beneficial ^59^ in this context. As WCMES increases cellular metabolism in fungal hyphae as it does with human cells, the pace of vesicle circulation and production would surpass the speed of hyphal extension, which operates at maximal speed as it always is the restrictive factor (as proven by the very dynamics of apical branching). Thus, in such cases only lateral branching can absorb the increased production of hyphal growth elements. In multicellular fungi, the vesicle immigration/traffic and accumulation are similar functions to the abovementioned and lead to analogous results; it has been established that a hyphal tip extends when it is being well-supplied in vesicles, which accumulate by it preferentially (apical dominance) ^60,61^. Once vesicle supply increases over a threshold, i.e. if the extension rate of the hypha slows down, biases to keep the vesicle influx accumulated are overrun and apical branching occurs. If, on the other hand, the flow of vesicles along the axis of the hyphal cells, through septal pores (where available) is compromised, a new tip (and thus branch) emerges at any new vesicle accumulation area ^55,62^. The relaxation of apical dominance because of protoplasmic flow fallout due to distance is a key factor in the occurrence of random lateral branching. Septum-related lateral branching, on the other hand, is triggered by new septum lockdown and pause of protoplasmic flow due to pore lockdown. Both random and septum-related branching occur at sites where vesicles accumulate due to different causes of compromising vesicle transition and orientation.

## Conclusions

The folding of the mycelial mat even in single-CFU mycelia produced from open-dish passive air sampling implies disproportional lateral and axial growth rates. As split-tip branching tends to maintain the angular density of hyphae steady, backed by fusion events and change of branching rate whenever necessary ^47^, the folds imply a spatially disproportionate arrangement, obviously a lateral expansion absorbed into a 3rd dimension arrangement. This arrangement may be *ad hoc* and asymmetric or very tedious and symmetric. The latter implies a smoother and more even succession and climax of events. The farther edge of such radial and symmetrical folds forming a ring may signify either a transition to preferentially apical/axial growth and branching after a phase of predominantly-or simply intense-lateral growth; or transition between different phases of hyphal growth ratios, most probably from exponential to transitional to linear. Successive rings of folds might imply repetition of such transitional phenomena, the duration perhaps deducible by the radial length of the folds. This incremental growth, seem in here as an effected result but also observed without ES treatment ^39^ might have ramifications in elucidating and understanding infection dynamics of mycelial fungi, and in the longer term, allow some innovative treatment options.

## Acknowledgements

As per ICMJE prerogatives the authors wish to thank Ass Prof Kostantinos Poulas, Dept of Pharmacy, University of Patras, for lending the Wetling W200 device to the HCPF/UoA.

## Authors Contributions

As per the CRediT (Contributor Roles Taxonomy), Manousos E. Kambouris conceived the operative idea, developed the peculiar aspects of methodology and produced the manuscript (Writing-Original Draft). Katerina Karageorgou and Gian-Marco Ludovici contributed in the research and investigation, the respective processing and the medical and biological interpretation (respectively) of the results, as well as in both phases of writing (Original Draft and Review and Editing. Stavroula Kritikou and Afroditi Milioni engaged in Methodology, results validation and interpretation and in drafting the original manuscript. All authors made a significant intellectual contribution, read and approved the final version. Aristea Velegraki undertook the supervision of the project, participated in experiments (Methodology) and results validation and interpretation, and in the editing process (Writing - Review & Editing).

